## Supplementary material for "Inhibition of mitochondrial protein import and proteostasis by a pro-apoptotic lipid": Supplemetal Figure 1

### Slide 1
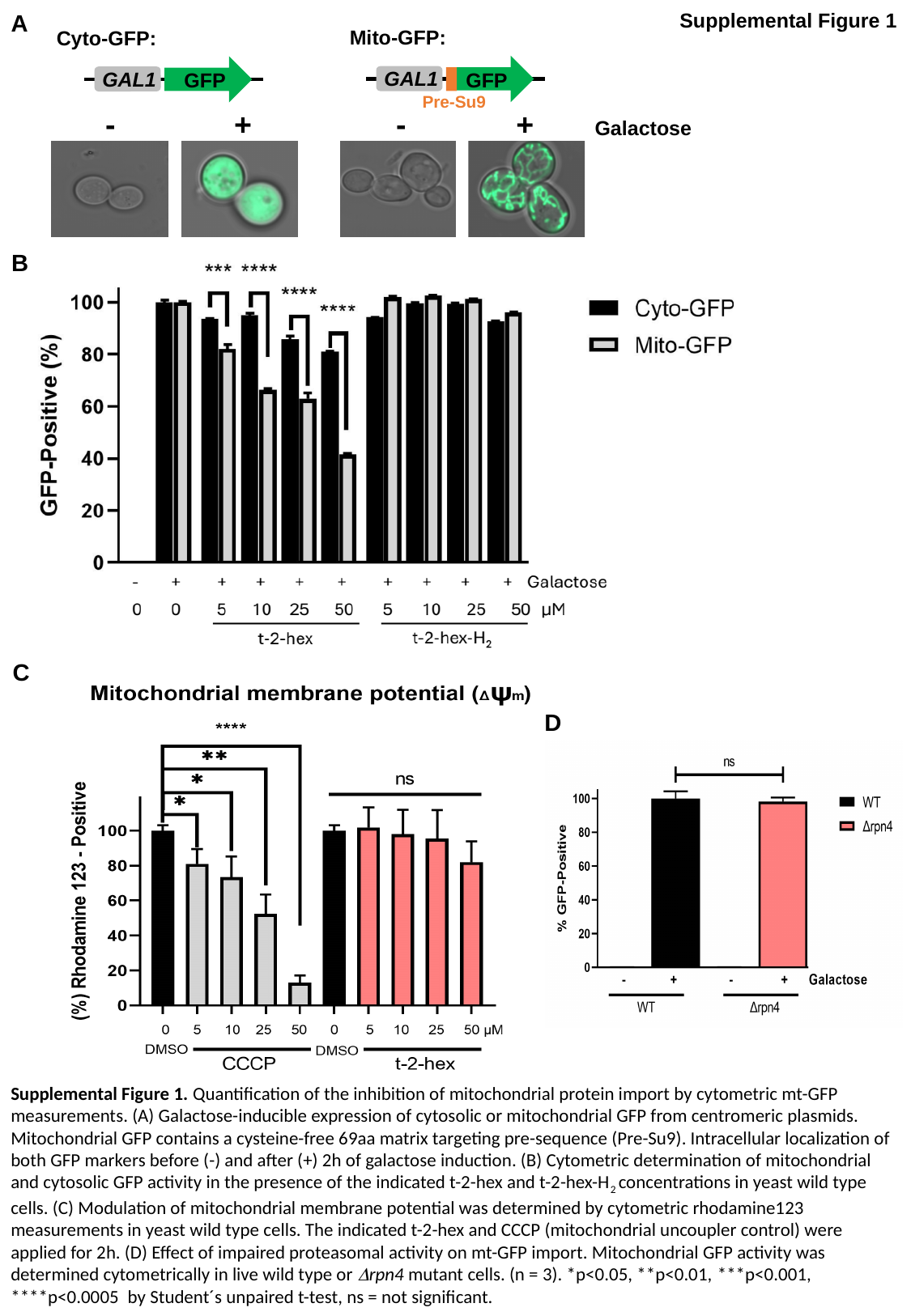

Supplemental Figure 1
A
Mito-GFP:
Cyto-GFP:
GAL1
GFP
GAL1
GFP
Pre-Su9
- + - + Galactose
B
C
D
Supplemental Figure 1. Quantification of the inhibition of mitochondrial protein import by cytometric mt-GFP measurements. (A) Galactose-inducible expression of cytosolic or mitochondrial GFP from centromeric plasmids. Mitochondrial GFP contains a cysteine-free 69aa matrix targeting pre-sequence (Pre-Su9). Intracellular localization of both GFP markers before (-) and after (+) 2h of galactose induction. (B) Cytometric determination of mitochondrial and cytosolic GFP activity in the presence of the indicated t-2-hex and t-2-hex-H_2_ concentrations in yeast wild type cells. (C) Modulation of mitochondrial membrane potential was determined by cytometric rhodamine123 measurements in yeast wild type cells. The indicated t-2-hex and CCCP (mitochondrial uncoupler control) were applied for 2h. (D) Effect of impaired proteasomal activity on mt-GFP import. Mitochondrial GFP activity was determined cytometrically in live wild type or Δrpn4 mutant cells. (n = 3). *p<0.05, **p<0.01, ***p<0.001, ****p<0.0005 by Student´s unpaired t-test, ns = not significant.
