## Supplemental Figure 2 for "Inhibition of mitochondrial protein import and proteostasis by a pro-apoptotic lipid"

### Slide 1
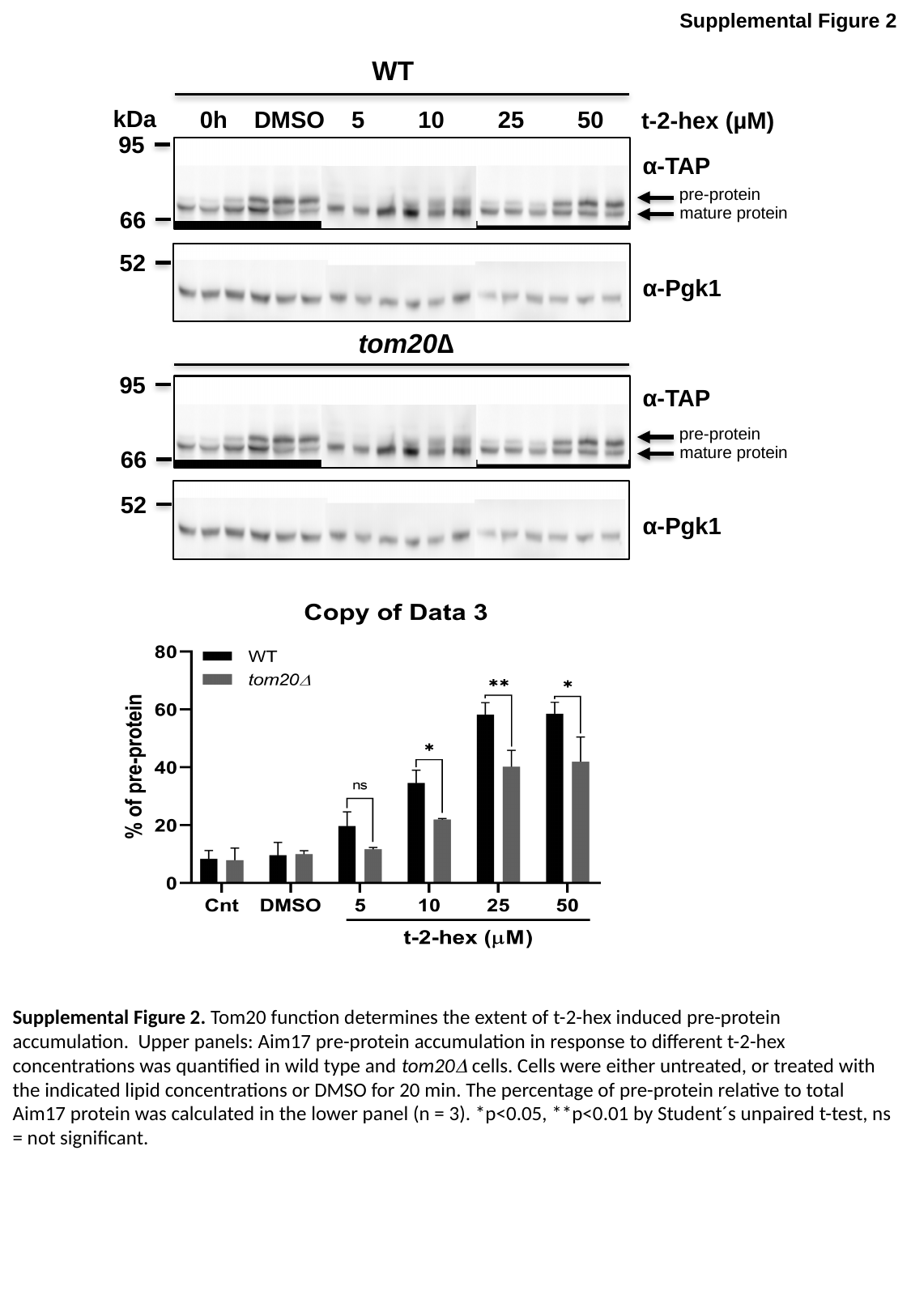

Supplemental Figure 2
WT
kDa
 0h DMSO 5 10 25 50
t-2-hex (µM)
95
α-TAP
pre-protein
mature protein
66
52
α-Pgk1
tom20∆
95
α-TAP
pre-protein
mature protein
66
52
α-Pgk1
Supplemental Figure 2. Tom20 function determines the extent of t-2-hex induced pre-protein accumulation. Upper panels: Aim17 pre-protein accumulation in response to different t-2-hex concentrations was quantified in wild type and tom20Δ cells. Cells were either untreated, or treated with the indicated lipid concentrations or DMSO for 20 min. The percentage of pre-protein relative to total Aim17 protein was calculated in the lower panel (n = 3). *p<0.05, **p<0.01 by Student´s unpaired t-test, ns = not significant.
